# MYC2 mediated regulation of xylan substitution patterns

**DOI:** 10.64898/2026.06.11.731540

**Authors:** Shaogan Wang, Markus Pauly, Vicente Ramírez

## Abstract

*O-*Acetylation is the most abundant xylan decoration in eudicot plants and plays a critical role in determining xylan conformation and its interactions with cellulose and lignin, thereby contributing to secondary cell wall (SCW) integrity. In *Arabidopsis*, loss of the xylan *O*-acetyltransferase TBL29/ESK1 causes collapsed xylem and growth defects that can be suppressed by mutations in strigolactone (SL) biosynthesis genes such as *MAX3*. However, the molecular basis of this suppression remains unknown.

Hypoacetylated xylan in *tbl29* has a higher frequency of methyl glucuronic acid (MeGlcA) substituents, while the ratio of GlcA/MeGlcA is recovered in *tbl29 max3*. Furthermore, gene expression analyses reveal that the three xylan glucuronoxylan methyltransferases (GXM1/2/3) involved in xylan MeGlcA modification are upregulated in *tbl29* SCWs but downregulated in *tbl29 max3*. Genetic analysis shows that the transcription factor MYC2 is required for *max3*-mediated suppression: the loss of MYC2 in *tbl29 max3* prevents growth recovery and reverts *GXM* genes expression and xylan MeGlcA substitution levels. We propose a model where SL deficiency enhances *MYC2* transcription, which in turn represses *GXMs*, thereby fine-tuning xylan methylation and re-establishing the MeGlcA/GlcA substitution balance under conditions of reduced *O*-acetylation.

Our findings identify a MYC2-dependent regulatory module linking SL signalling to xylan methylation and reveal a genetically encoded compensatory mechanism that mitigates the consequences of defective xylan *O*-acetylation. More broadly, this work demonstrates that plants can preserve SCW function through adaptive remodelling of polysaccharide substitution patterns, highlighting an unexpected plasticity in SCW biosynthesis.

**Significance Statement:** Secondary cell wall integrity depends on the coordinated modification of xylan. We show that defects caused by reduced xylan *O*-acetylation can be alleviated through a strigolactone- and MYC2-dependent pathway that alters xylan methylglucuronidation. Rather than restoring the original wall composition, this mechanism appears to compensate for the loss of *O*-acetyl groups by remodelling polysaccharide substitution patterns to maintain cell wall function, revealing a new layer of plasticity in secondary wall biosynthesis.

## INTRODUCTION

Plant cell walls are dynamic, polysaccharide-rich composites that play essential roles in plant growth, development, and adaptation to environmental stresses (Zhang et al., 2021; Delmer et al., 2024). The hemicellulose xylan is one of the major structural components of plant secondary cell walls (SCW), where it interacts with cellulose and lignin to build a rigid yet partially flexible network (Simmons et al., 2016; Martinez-Abad et al., 2017; Kang et al., 2019; Zhong et al., 2019). Xylan consists of a linear backbone of β-(1→4)-linked xylosyl residues that can be decorated with various substituents (Scheller and Ulvskov, 2010; Pauly et al., 2013; Ye and Zhong, 2022; Qaseem et al., 2025). In dicotyledonous plants such as the model species *Arabidopsis thaliana*, xylosyl residues are commonly modified with acetyl groups at the *O*-2 and/or *O*-3 positions (Gille and Pauly, 2012; Pauly and Ramirez, 2018). These modifications are mediated by Xylan *O*-acetyltransferases from the Trichome Birefringence-Like (TBL) family such as TBL29/ESKIMO1/XOAT1 (Lee et al., 2011; Manabe et al., 2013; Yuan et al., 2013; Schultink et al., 2015; Yuan et al., 2016a; Yuan et al., 2016b).

In addition to *O*-acetylation, xylan can be substituted with single α-D-glucuronic acid (GlcA) residues at the *O*-2 position by glucuronyltransferases from the glycosyl transferase (GT) 8 family (Mortimer et al., 2010; Lee et al., 2012a). In *Arabidopsis*, GLUCURONIC ACID SUBSTITUTION OF XYLAN (GUX) enzymes decorate different domains of the xylan backbone in SCWs (Bromley et al., 2013). GlcA residues can be further methylated to form 4-*O*-methyl-α-D-glucuronic acid (MeGlcA), a reaction catalyzed by GLUCURONOXYLAN METHYLTRANSFERASE1 (GXM1), GXM2, and/or GXM3/AtGXMT1 (Urbanowicz et al., 2012; Lee et al., 2012b; Yuan et al., 2014). Together, these modifications contribute to the structural diversity and functional complexity of xylan.

Genetic studies have revealed that different xylan substitutions contribute unequally to SCW function. Arabidopsis mutants in which xylan is largely devoid of GlcA or MeGlcA substitutions such as in the Arabidopsis *gux1gux2* double mutant or the *gxm1/2/3* triple mutant display relatively mild phenotypes (Mortimer et al., 2010; Lee et al., 2012a). In contrast, alterations in xylan *O*-acetylation have severe consequences for SCW integrity. For example, the *tbl29* mutant, containing 43% less *O*-acetyl substituents on its xylan, exhibits collapsed xylem vessels, reduced growth, and altered responses to multiple biotic and abiotic stresses, a collection of pleiotropic phenotypes collectively referred to as the xylan hypoacetylation syndrome (Lee et al., 2011; Xiong et al., 2013; Schultink et al., 2015; Gao et al., 2017). Similar syndromes have been described in Arabidopsis, rice or poplar, supporting the hypothesis that hypoacetylation renders xylan dysfunctional, altering its interactions with cellulose and lignin, thereby weakening SCWs in vascular tissues and compromising their ability to withstand the mechanical demands associated with water transport (Grantham et al., 2017; Joshi and Gupta, 2026).

Despite the severe consequences of a dysfunctional xylan, the mechanisms specifically regulating xylan modifications, particularly xylan *O*-acetylation, remain poorly understood. Specific transcriptional regulators have not been identified, but TBL, GUX and GXM expression profiles suggest co-regulation with secondary wall biosynthetic programs and are likely integrated within NAC-MYB regulatory networks controlling SCW formation (Didi et al., 2015; Nakano et al., 2015; Zhong and Ye, 2015; Im and Son, 2025). MYC transcription factors, particularly MYC2 and its paralogue MYC4, have been shown to transcriptionally regulate specific xylan *O*-acetyltransferase genes, including TBL37, in a jasmonate-dependent manner, suggesting a potential link between stress signalling and cell wall polysaccharide *O*-acetylation (Sun et al., 2020). However, whether MYC-dependent transcriptional changes translate into altered xylan *O*-acetylation levels remains unclear.

Previous work has shown that overexpression of *GUX1* under the control of the *TBL29* promoter rescues the developmental defects of *tbl29* without restoring xylan *O*-acetylation levels (Xiong et al., 2015). This finding indicates that enhanced glucuronidation can compensate for reduced *O*-acetylation in vivo. In addition, hypoacetylated xylan in *tbl29* contains elevated levels of MeGlcA substitution, suggesting compensatory cross-talk between distinct xylan modifications (Grantham et al., 2017). However, the molecular basis of this response *i*.*e*., how xylan hypoacetylation leads to increased MeGlcA substitution, and whether xylan modification patterns are coordinately regulated, remains unknown.

Suppressor screenings have identified genetic factors that alleviate the hypoacetylation syndrome in *tbl29* without restoring *O*-acetylation levels, revealing the existence of additional genetically encoded compensatory pathways (Bensussan et al., 2015). For example, mutations in MORE AXILLARY BRANCHES 3 (MAX3) and MAX4, two genes required for the biosynthesis of the hormone strigolactone (SL), suppress *tbl29*’s dwarfism and enhanced stress tolerance. Exogenous application of synthetic SLs reverses this suppression, indicating that altered SL signalling is responsible (Ramirez et al., 2018; Ramirez and Pauly, 2019). However, the mechanistic basis for how SL deficiency compensates for xylan hypoacetylation has remained elusive.

Here, we establish that suppression of the *tbl29* hypoacetylation syndrome in *tbl29 max3* is mediated by altered MeGlcA substitution of xylan through MYC2-dependent transcriptional upregulation of GXM genes. By integrating genetic, transcriptomic and cell wall compositional analyses, we propose a model for a regulatory pathway linking SL signalling to xylan modification through MYC2, demonstrating that plants can employ transcriptional reprogramming of cell wall biosynthesis as a compensatory mechanism to maintain SCW integrity under hypoacetylation stress. Our findings establish a molecular framework for understanding how hormonal signals coordinate cell wall polymer modifications to ensure functional redundancy and developmental robustness.

## RESULTS

### Suppression of *tbl29*-associated growth defects by *max3* requires *GUX1*

Disrupting strigolactone (SL) biosynthesis in the *tbl29* mutant suppresses the developmental and stress-related defects caused by xylan hypoacetylation without restoring xylan *O*-acetylation levels suggesting the activation of a compensatory mechanism (Ramirez et al., 2018; Ramirez and Pauly, 2019). Since upregulation of *GUX1* has been shown to rescue the hypoacetylation syndrome in *tbl29* (Xiong et al., 2015), we investigated the effect of introducing a *gux1* mutation on *tbl29 max3*. The phenotype of the resulting *tbl29 max3 gux1* triple mutant and related genotypes was evaluated (Figure 1).

**Figure 1.**
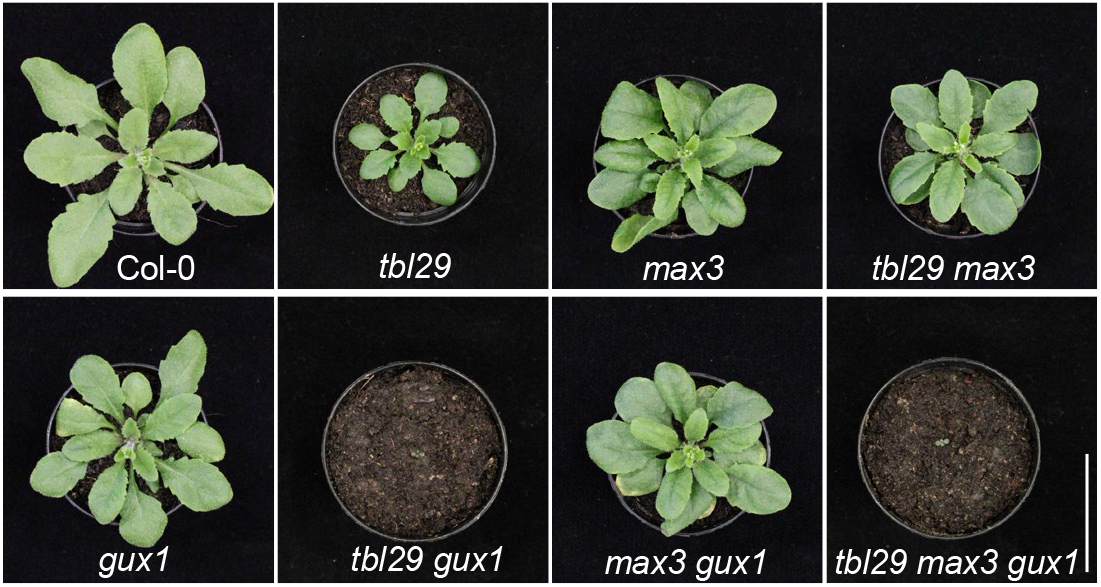
GUX1 is essential for *max3*-dependent suppression of the *tbl29* phenotypes. Representative images of 4-week-old plants of the indicated phenotypes. Bar = 5 cm.

As previously reported, while *tbl29* displayed reduced growth and smaller organs, the *gux1* mutant line showed near wild-type development. The *tbl29 gux1* double mutant was severely dwarfed, indicating that impaired xylan glucuronidation has additive detrimental effects in the hypoacetylated *tbl29* background. At the 4-week growth stage, the *tbl29 max3* suppressor already exhibited a clear recovery of the reduced leaf-size phenotype observed in *tbl29*. In contrast, *tbl29 max3 gux1* triple mutant plants displayed severe growth defects similar to *tbl29 gux1*, arresting development approximately 20 days after germination. These results indicate that GUX1 is required for the suppression of the xylan hypoacetylation syndrome in *tbl29 max3*.

### Suppression of *tbl29*-associated defects correlates with xylan methylation frequency

To assess whether alterations in xylan glucuronidation contribute to the phenotypic rescue observed in *tbl29 max3*, we quantified the absolute levels of xylose, methylglucuronic acid (MeGlcA), and glucuronic acid (GlcA) in stem cell wall material enriched in SCWs (Table 1). Throughout this study, the combined content of MeGlcA and GlcA is referred to as [Me]GlcA (Mortimer et al., 2010). The [Me]GlcA/Xyl molar ratio was calculated as an indicator of xylan substitution frequency. Both *tbl29* and *tbl29 max3* exhibited [Me]GlcA/Xyl ratios comparable to wild type indicating a similar degree of overall xylan [Me]GlcA substitution (Figure 2a). As controls, the *gux1* and *max3 gux1* mutants displayed a markedly reduced [Me]GlcA/Xyl ratio resulting from a reduced glucuronidation. Conversely, *GUX1* overexpression in the *tbl29* mutant (*tbl29:GUX1-OE* line) significantly increased the GlcA level consistent with previous findings (Xiong et al., 2015), resulting in an increased [Me]GlcA/Xyl ratio (Figure 2a).

**Table 1.** Absolute quantification of xylose, MeGlcA, and GlcA contents in 6-week-old stems.

| Genotype | Xyl | MeGlcA | GlcA | [Me]GlcA |
| --- | --- | --- | --- | --- |
| Col-0 | 132.30±9.78<sup>ab</sup> | 3.08±0.12<sup>ef</sup> | 3.03±0.43<sup>b</sup> | 6.12±0.48<sup>ce</sup> |
| *tbl29* | 139.65±12.47<sup>ab</sup> | 4.76±0.29<sup>ab</sup> | 2.02±0.12<sup>c</sup> | 6.78±0.34<sup>bcd</sup> |
| *max3* | 119.12±18.89<sup>b</sup> | 3.30±0.26<sup>de</sup> | 2.06±0.53<sup>c</sup> | 5.36±0.67<sup>e</sup> |
| *tbl29 max3* | 134.27±13.14<sup>ab</sup> | 4.00±0.16<sup>cd</sup> | 3.15±0.36<sup>b</sup> | 7.15±0.47<sup>bc</sup> |
| *gux1* | 155.56±0.16<sup>a</sup> | 2.40±0.29<sup>fg</sup> | 1.09±0.04<sup>d</sup> | 3.49±0.30<sup>f</sup> |
| *max3 gux1* | 135.03±0.16<sup>ab</sup> | 2.08±0.08<sup>g</sup> | 0.97±0.08<sup>d</sup> | 3.05±0.14<sup>f</sup> |
| *tbl29:GUX1-OE* | 144.07±0.16<sup>ab</sup> | 3.35±0.39<sup>de</sup> | 5.78±0.44<sup>a</sup> | 9.13±0.50<sup>a</sup> |
| *myc2* | 121.78±15.85<sup>b</sup> | 3.04±0.45<sup>ef</sup> | 2.72±0.07<sup>bc</sup> | 5.76±0.52<sup>de</sup> |
| *tbl29 myc2* | 147.79±13.48<sup>ab</sup> | 5.27±0.20<sup>a</sup> | 2.20±0.28<sup>c</sup> | 7.47±0.45<sup>bcd</sup> |
| *max3 myc2* | 127.06±7.34<sup>ab</sup> | 3.33±0.32<sup>de</sup> | 2.05±0.03<sup>c</sup> | 5.38±0.34<sup>b</sup> |
| *tbl29 max3 myc2* | 135.35±12.46<sup>ab</sup> | 4.53±0.41<sup>bc</sup> | 2.04±0.28<sup>c</sup> | 6.57±0.40<sup>e</sup> |

Values represent means ± SD from four biological replicates. Significant differences within each genotype were determined by one-way ANOVA followed by Tukey's multiple comparison test, and are indicated using a compact letter display (letters in common indicate no significant difference, $P > 0.05$ ).

**Figure 2.**
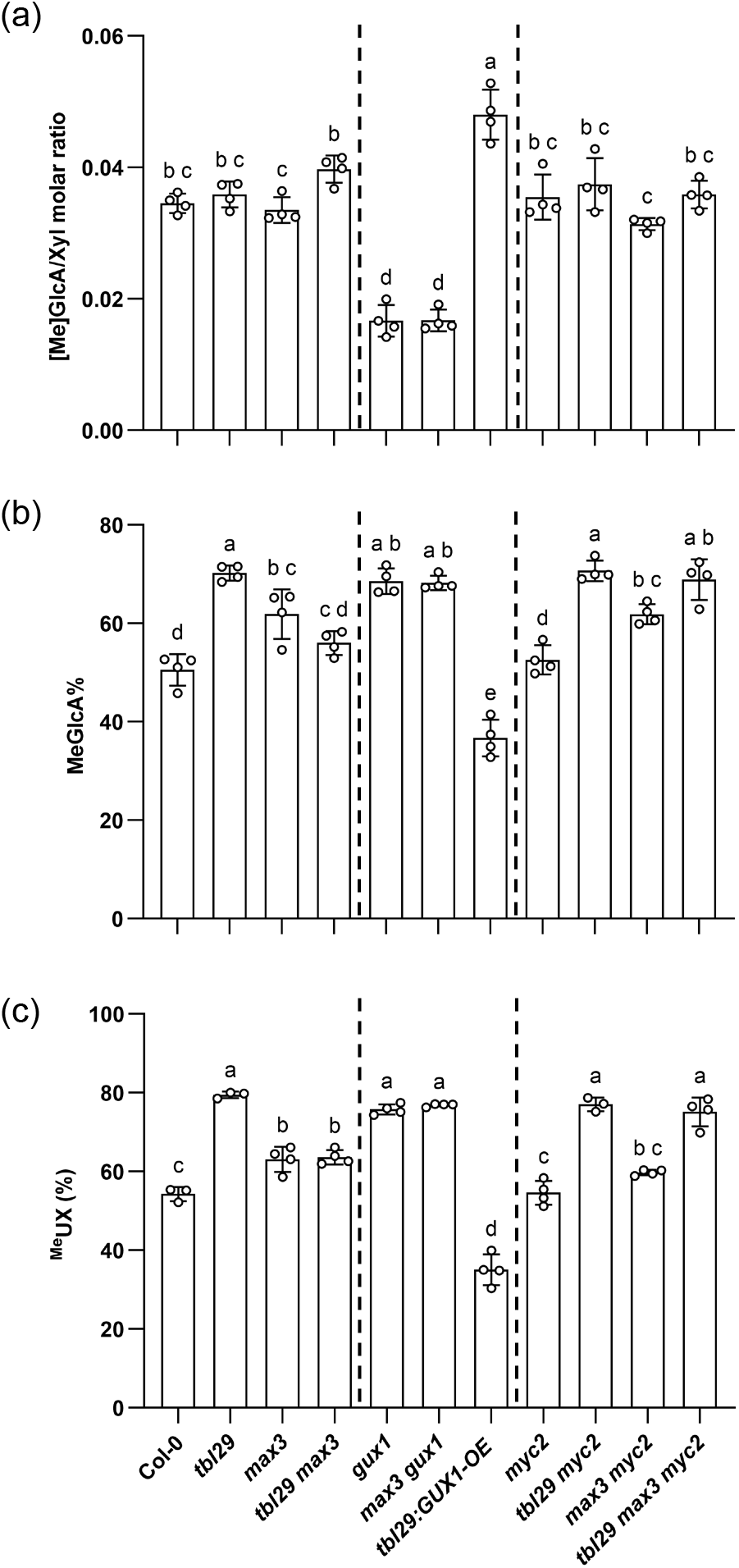
Xylan GlcA and MeGlcA substitution pattern in *tbl29*-related genotypes. (a)Degree of overall xylan glucuronidation. The [Me]GlcA/Xyl molar ratio was calculated based on the absolute amounts of [Me]GlcA (methylated and unmethylated GlcA) and xylose released from TFA-hydrolyzed AIR of 6-week-old stem. (b)Degree of xylan MeGlcA substitution. The percentage of MeGlcA (MeGlcA%) represents the proportion of methylated GlcA relative to the total GlcA pool ([Me]GlcA). (c)Relative abundance of MeGlcA-substituted xylooligosaccharides. Xylooligosaccharides were released from deacetylated stem AIR by GH10 xylanase digestion and analyzed by MALDI-TOF mass spectrometry. Full spectra are shown in Figure S1. Relative abundances were calculated based on the intensities of the corresponding ion signals. U, GlcA; X, xylose; Me, methyl. Data are represented as means ± SD (n ≥3 biological replicates). Different letters indicate significant differences (Tukey’s HSD test, p<0.05).

The comparable [Me]GlcA levels between the *tbl29* mutant and *tbl29 max3* suppressor suggest that *max3*-dependent suppression of *tbl29*-associated defects is unlikely to result from a compensatory effect of increased xylan glucuronidation. However, differences in the proportion of MeGlcA relative to the total [Me]GlcA pool (MeGlcA%) were observed. As previously reported, the proportion of methylated GlcA was significantly increased in the *tbl29* mutant compared to wild type (Figure 2b). Notably, the *tbl29 max3* suppressor displayed a significant reduction in MeGlcA% relative to *tbl29*, comparable with wild-type levels. Consistent with a reduced pool of non-methylated GlcA residues, the *gux1* and *max3 gux1* mutants exhibited a strong increase in MeGlcA%. In contrast, *tbl29:GUX1-OE* plants exhibited a marked decrease in MeGlcA% in accordance with an increased abundance of non-methylated GlcA.

To further validate the observed alterations in xylan methylglucuronidation associated to the phenotypic rescue in *tbl29 max3*, we performed xylan oligosaccharide mass profiling (OLIMP) and matrix-assisted laser desorption/ionization time-of-flight (MALDI-ToF) mass spectrometry analysis. Alcohol-insoluble residue (AIR) samples were first deacetylated by alkaline treatment and subsequently digested with a GH10 endoxylanase. This treatment releases a series of xylooligosaccharides which are diagnostic of the xylan substitution pattern. Inspection of the resulting mass spectra revealed clear differences in relative abundance of MeGlcA -substituted (^Me^UX) versus GlcA-substituted (UX) xylooligosaccharides (Figure S1). Semi-quantitative analysis of the ion signals revealed an increased 79% ^Me^UX relative abundance (^Me^UX%) in *tbl29* compared to 54% in wild type, indicative of a higher frequency of MeGlcA decorations (Figure 2c). In contrast, the *tbl29max3* suppressor displayed reduced ^Me^UX% levels of approximately 64% compared with the *tbl29* mutant similar to the *max3* parental line. Consistent with the differences observed in the [Me]GlcA/Xyl molar ratio and MeGlcA%, the ^Me^UX% was increased to approximately 76% in the *gux1* and *max3 gux1* mutants, and decreased to approximately 35% in the *tbl29:GUX1-OE* transgenic plants (Figure 2c).

In summary, these results indicate that altered overall xylan glucuronidation does not seem to be the structural basis for *max3*-dependent suppression of *tbl29*-associated defects. Instead, the suppression is specifically associated with a downregulation of xylan methylglucuronidation.

In *Arabidopsis*, three glucuronoxylan methyltransferases, GXM1, GXM2, and GXM3/AtGXMT1, have been reported to catalyze the transfer of methyl groups onto GlcA side chains of xylan. To determine whether the restoration of the degree of MeGlcA substitutions observed in *tbl29 max3* is associated to altered *GXM* gene expression, we determined transcript abundance by quantitative RT-PCR (qPCR) (Figure 3). In line with the altered MeGlcA/GlcA substitution pattern identified by composition and xylan mass profiling analyses (Figures 2b,c), qPCR revealed that *GXM1, GXM2*, and *GXM3* were strongly upregulated in the *tbl29* mutant suggesting a contribution to the increase in xylan methylglucuronidation. In the *tbl29 max3* suppressor, their expression was significantly downregulated. These results further support that the phenotypic rescue observed in *tbl29 max3* is associated to a restoration of wild type xylan methylglucuronidation through the downregulation of *GXMs* expression.

**Figure 3.**
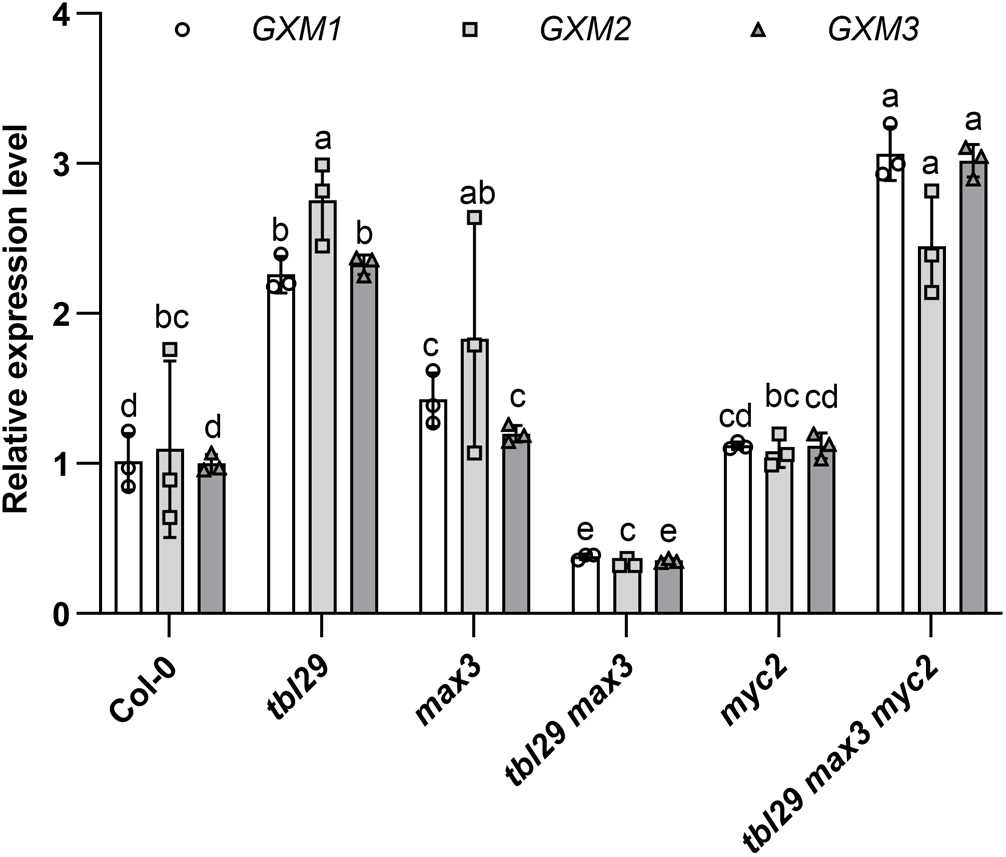
Transcriptional regulation of *GXM* genes associated to altered degree of xylan MeGlcA. Quantitative real-time PCR analysis of *GXM1, GXM2*, and *GXM3* gene expression in stem tissue. Values were normalized to *AtACTIN2* gene expression. Data are represented as means ± SD (n = 3 biological replicates). Different letters indicate significant differences (Tukey’s HSD test, p<0.05).

### MYC2 is required for *max3*-dependent suppression of *tbl29*-associated growth defects

In light of the altered *GXMs* expression patterns, we sought to identify components of the transcriptional regulatory program involved. For this purpose, we analyzed the 2-kb upstream promoter regions of the *GXMs* to search for conserved transcription factor binding motifs by PlantCARE database (http://bioinformatics.psb.ugent.be/webtools/plantcare/html/) (Lescot et al., 2002). This analysis identified cis-elements common to all three *GXM* promoter regions predicted to bind bZIP, MYB, and MYC transcriptional factors (Table S1). These include the well characterized CANNTG recognition motif for MYC2 (Li et al., 2014; Lopez-Vidriero et al., 2021), a key transcription factor in the jasmonate signaling pathway, previously implicated in SCW-associated transcriptional programs (Sun et al., 2020; Luo et al., 2022; Im et al., 2024). Interestingly, silencing of SL biosynthesis genes such as *MAX3, MAX4*, or *MAX1* resulted in increased transcription of *MYC2* in tomato roots (Xu et al., 2019). Together, these observations suggested that MYC2 could function as a candidate mediator linking SL deficiency to *GXM* genes regulation.

To determine whether MYC2 is involved in *max3*-dependent suppression of *tbl29*, we introduced a well characterized *myc2* mutation (Anderson et al., 2004; Jung et al., 2015; Zhang et al., 2023) into the *tbl29 max3* background to generate a *tbl29 max3 myc2* triple mutant. Detailed phenotypic analyses were performed for all corresponding genotypes (Figure 4). At the mature stage, *myc2* plants exhibited a wild-type stature and architecture, whereas *tbl29 myc2* resembled *tbl29*, displaying dwarfism and altered shoot architecture producing on average half the number of shoot branches observed in wild-type plants (Figure 4a-c). The *tbl29 max3 myc2* triple mutant displayed reduced leaf size and dwarfism comparable to *tbl29* and *tbl29 myc2* mutants, indicating that the rescued phenotype observed in the *tbl29 max3* mutant combination is abolished in the absence of MYC2.

**Figure 4.**
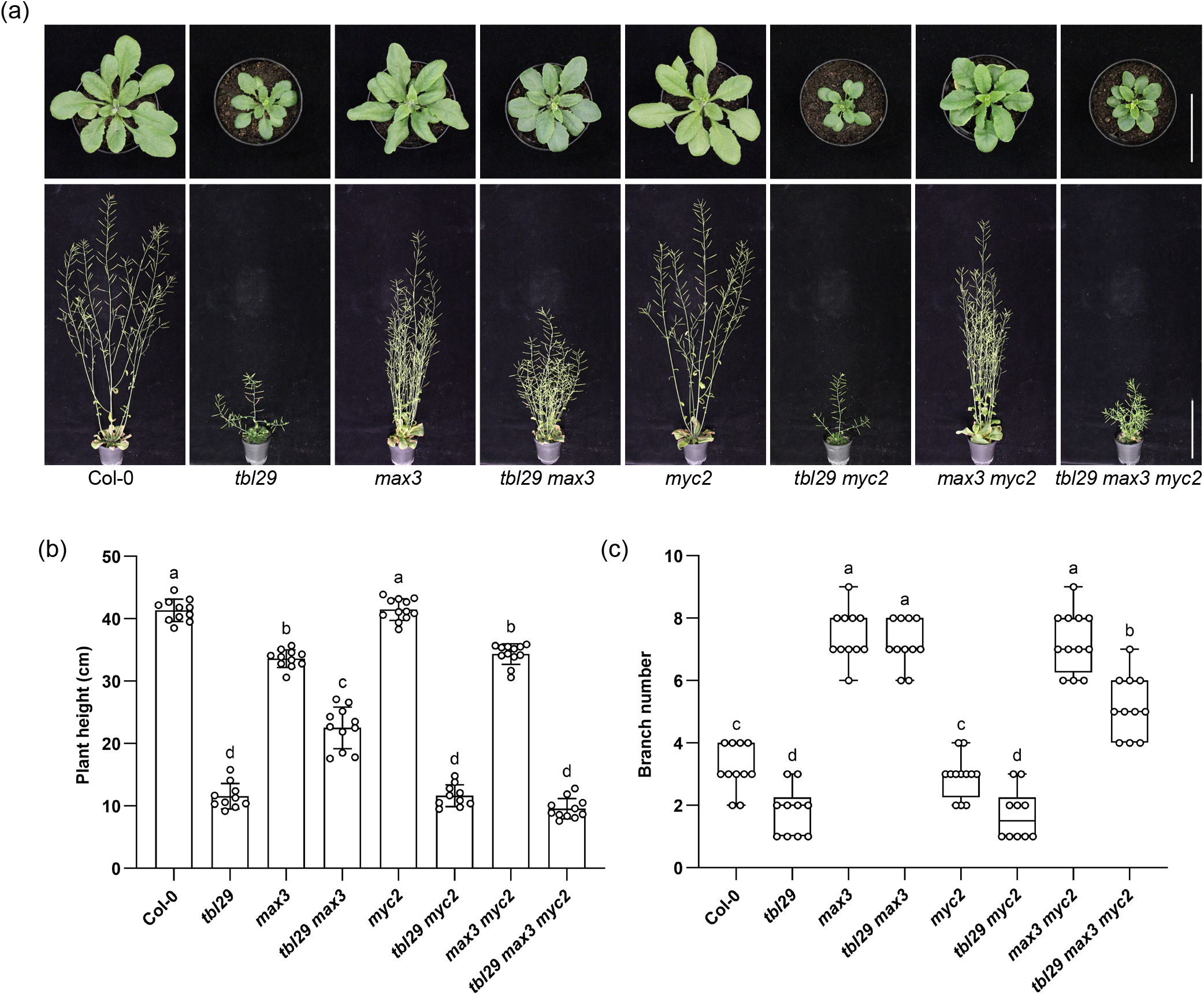
MYC2 is required for *max3*-dependent suppression of the *tbl29* phenotypes. (a)Growth phenotypes of 4-week-old (upper panel) and 6-week-old (lower panel) plants. Scale bars = 5 cm (upper panel) and 10 cm (lower panel). (b)Measurements of primary inflorescence stem height. Data are represented as means ± SD. (c)Number of axillary branches developed after 6 weeks. The box boundaries represent the 25th (lower), 50th (median), and 75th (upper) percentiles. Whiskers extend to the absolute minimum and maximum values of the dataset. The internal solid line indicates the mean. White circles represent individual data points (spot distances). Different letters in b and c indicate significant differences (Tukey’s HSD test, p < 0.05, n ≥10 biological replicates).

The phenotypic recovery in *tbl29 max3* included not only a significant increase in plant height compared with the *tbl29* mutant, but also a characteristic increased number of shoot branches typical of SL-deficiency (Sorefan et al., 2003; Booker et al., 2004; Figure 4c). The increased number of shoot branches was observed in all mutant combinations including *max3* such as *max3, max3 myc2*, and *tbl29 max3 myc2* indicating that MYC2 seems to be required for phenotypes specifically associated to xylan hypoacetylation and not to other SL-controlled processes such as axillary branch formation.

Overall, our genetic evidence strongly supports the notion that *max3*-dependent suppression of the *tbl29* hypoacetylation syndrome requires the MYC2 transcriptional regulator.

### Altered xylan methylation and *GXMs* gene expression in *tbl29 max3* are MYC2-dependent

As genetic evidence indicated that MYC2 is required for suppression of the *tbl29*-associated defects, we next examined xylan GlcA and MeGlcA substitution levels in mature stems of the various *myc2* mutant combinations (Figure 2). Quantification of xylose, MeGlcA, and GlcA contents (Table 1) revealed that the *myc2* single mutant displayed a [Me]GlcA/Xyl molar ratio comparable to wild type. Similar results were also observed when comparing the *max3 myc2* to *max3* mutants (Figure 2a), indicating that loss of MYC2 function does not affect xylan glucuronidation. In contrast, the increased MeGlcA% observed in the phenotypically rescued *tbl29 max3* plants was abolished in the *tbl29 max3 myc2* triple mutant (Figure 2b).

To further validate these results, we also performed xylan OLIMP following GH10 hydrolysis (Figure S1). Spectra obtained from *tbl29 myc2* and *max3 myc2* resembled those of *tbl29* and *max3*, respectively. This result indicates that *myc2* does not influence the increased frequency of MeGlcA decorations observed in *tbl29*, consistent with the composition analysis shown in Figure 2b. In contrast, OLIMP profiles from *tbl29 max3 myc2* differed markedly from *tbl29 max3* and instead resembled *tbl29*. Semi-quantitative analyses further revealed increased relative ion abundance of MeGlcA-substituted xylooligosaccharides in *tbl29 max3 myc2* compared to *tbl29 max3* (Figure 2c). Collectively, these results demonstrate that MYC2 is required for the restoration of wild-type xylan methylglucuronidation during *max3*-dependent suppression of *tbl29*.

We next quantified *GXM1, GXM2*, and *GXM3* transcript abundance in stems of *myc2* and *tbl29 max3 myc2* plants and related mutant combinations (Figure 3). In *myc2, GXM* transcript levels were comparable to wild type, suggesting that loss of MYC2 alone does not influence *GXMs* expression under normal conditions. However, in *tbl29 max3 myc2*, expression of all three *GXM* genes was strongly upregulated relative to *tbl29 max3*, resembling the elevated transcript levels observed in *tbl29*.

Together, these results suggest that the reduction in xylan methylglucuronidation observed in the *tbl29 max3* suppressor is MYC2-dependent and correlates with altered *GXM* gene expression.

### SL deficiency promotes *MYC2* upregulation during suppression of *tbl29*-associated defects

To investigate how MYC2 mediates *max3* dependent suppression of *tbl29* at the molecular level, we examined *MYC2* transcript abundance in stems of Col-0, *tbl29, max3*, and *tbl29 max3* plants using qPCR (Figure 5). *MYC2* expression was comparable between *tbl29* and wild type indicating that xylan hypoacetylation alone does not alter *MYC2* transcription. In contrast, *MYC2* transcript levels were significantly elevated in *max3*, suggesting that, as already reported in tomato, SL deficiency promotes *MYC2* expression. In *tbl29 max3, MYC2* expression was lower than in *max3* but remained approximately two-fold higher than in *tbl29*. Collectively, these findings suggest that elevated *MYC2* expression resulting from SL deficiency might contribute to the compensatory mechanism underlying *max3*-dependent suppression of the *tbl29* xylan hypoacetylation syndrome.

**Figure 5.**
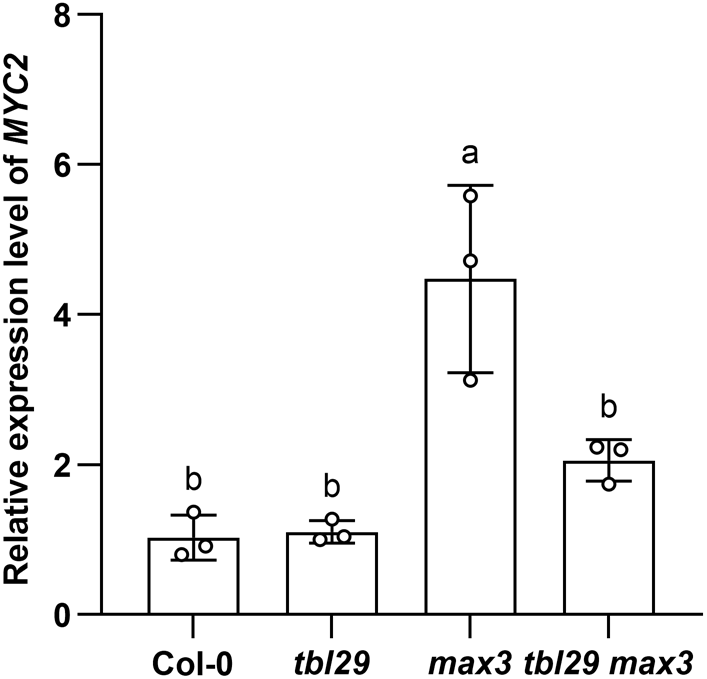
SL deficiency upregulates *MYC2* gene expression. Quantitative real-time PCR analysis of *MYC2* gene expression in stem tissue normalized to *AtACTIN2*. Data are represented as means ± SD (n = 3 biological replicates). Different letters indicate significant differences (Tukey’s HSD test, p<0.05).

## DISCUSSION

The structural integrity of the SCW depends on the coordinated structural modification of xylan, among which *O*-acetylation represents a major substitution that critically influences wall architecture and polymer interactions (Simmons et al., 2016; Kang et al., 2019; Kirui et al., 2022; Kundu et al., 2025). Loss of function of *TBL29*, the major xylan *O*-acetyltransferase, causes a reduction in *O*-acetylation which in turn deregulates other xylan modifications increasing the frequency of methylated GlcA substituents (Grantham et al., 2017; Figure 2b). It has been shown that mis-substituted xylan in *tbl29* fails to adopt a flattened ribbon-like twofold screw conformation required for the interaction with hydrophilic faces of cellulose fibrils (Grantham et al., 2017; Pfaff et al., 2024). As a result, reduced xylan *O*-acetylation in the *tbl29* mutant causes SCW abnormalities leading to severe growth defects such as dwarfism and collapsed xylem, accompanied by multiple alterations in responses to environmental cues (Xin and Browse, 1998; Bouchabke-Coussa et al., 2008; Lefebvre et al., 2011; Xiong et al., 2013; Escudero et al., 2017; Pauly and Ramirez, 2018).

Disruption of the biosynthesis of the hormone SL in the *tbl29* background can alleviate those SCW defects but without restoring xylan *O*-acetylation indicating that plants can activate compensatory mechanisms to buffer perturbations in xylan substitution patterns (Ramirez et al., 2018; Ramirez and Pauly, 2019; Figures 1 and 4). Here, we propose a model where SL deficiency partially compensates for defective xylan *O*-acetylation through a MYC2-dependent regulatory module that downregulates the expression of glucuronoxylan methyltransferase genes (*GXMs*), thereby reducing xylan MeGlcA content and restoring a more balanced MeGlcA/GlcA substitution pattern. Rather than re-establishing the original wall composition, this mechanism may enable the formation of an alternative polysaccharide substitution pattern that preserves wall functionality despite reduced *O*-acetylation (Figure 6).

**Figure 6.**
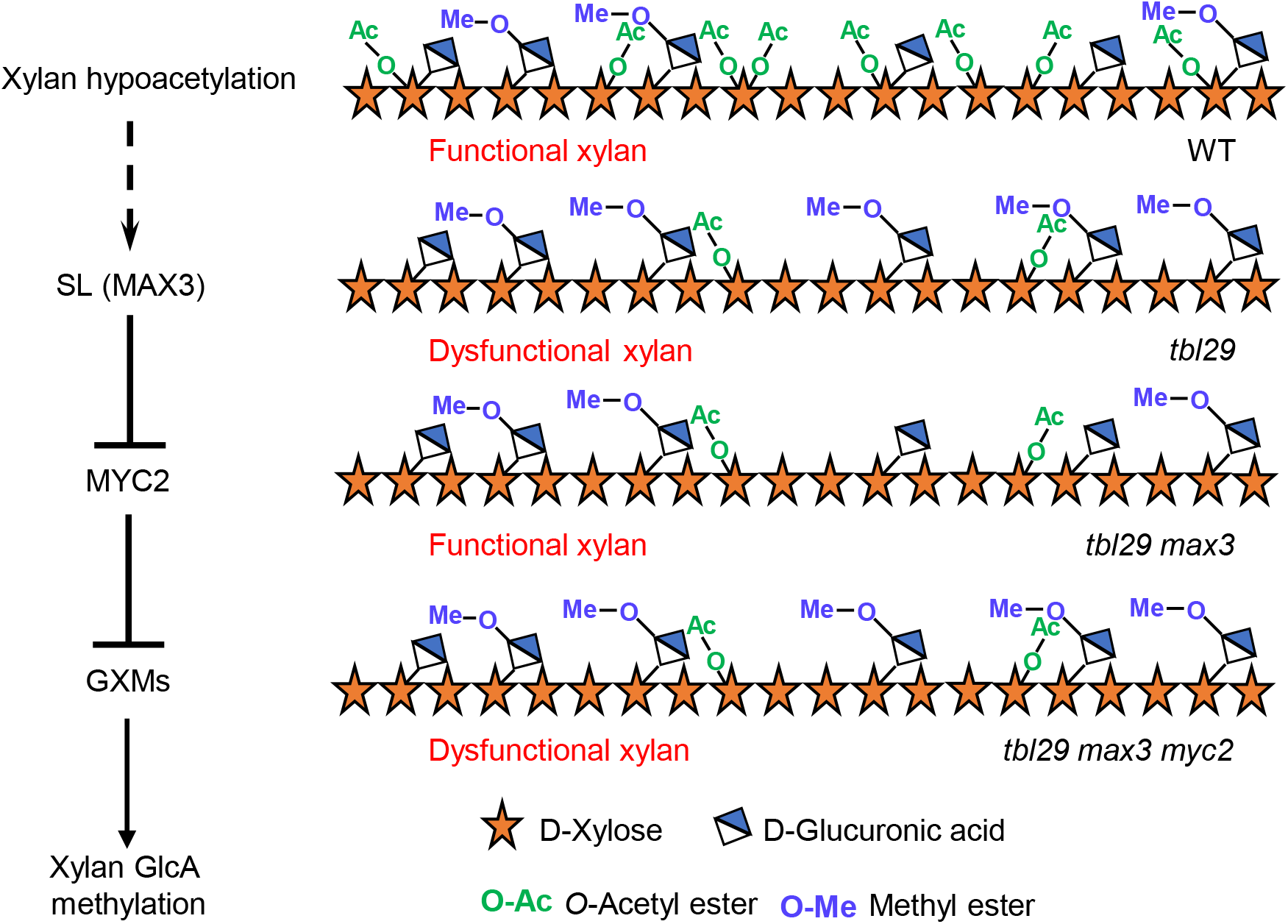
Proposed model of the SL-MYC2-GMXs module in the suppression of xylan hypoacetylation-associated growth defects. Xylan hypoacetylation in *tbl29* triggers overaccumulation of MeGlcA substituents resulting in disrupted secondary wall integrity. In the *tbl29 max3* suppressor, SL deficiency enhances MYC2 activity repressing *GXMs*, leading to reduced MeGlcA and restored xylan functionality. In contrast, loss of MYC2 derepresses *GXMs*, resulting in a xylan substitution pattern similar to *tbl29* and abolishing this compensatory effect.

Although xylan glucuronidation appears to be a structural prerequisite for *max3*-dependent suppression (Figure 1), the suppression mechanism does not rely on increasing the overall degree of glucuronidation. Total [Me]GlcA/Xyl ratio and *GUX1* expression remain unchanged in *tbl29 max3* relative to *tbl29* (Figures 2a and S2). Instead, the phenotypic recovery in *tbl29 max3* correlates with a downregulation of *GXM* gene expression reducing the frequency of methylation of GlcA substituents (MeGlcA%) (Figure 2b,c). Although we cannot exclude the possibility that these differences in the MeGlcA/GlcA substitution ratio partly reflect altered proportions of tissue compositions such as the abundance of fiber and vessel cells between the dwarf and recovered plants, the correlation between reduced *GXM* expression and reduced MeGlcA abundance supports the contribution of transcriptional regulation to this shift. Mechanistically, reducing MeGlcA content in xylan may compensate for reduced *O*-acetylation by altering polymer interactions. Methyl groups on GlcA add hydrophobic character and could interfere with hydrogen bonding or ionic interactions between xylan, cellulose, and/or lignin (Pereira et al., 2017; Kirui et al., 2022; Tryfona et al., 2023). Indeed, increased glucuronidation has been shown to rescue *tbl29* defects (Xiong et al., 2015) consistent with this compensatory mechanism. Thus, the restored MeGlcA/GlcA balance in the *tbl29 max3* suppressor may represent a compensatory remodeling of xylan substitution patterns that helps maintain functional interactions within the cellulose–xylan–lignin network despite reduced *O*-acetylation.

Several questions remain open. How does SL deficiency lead to elevated MYC2? One possibility is cross-talk with jasmonate (JA) signalling, as MYC2 is a master regulator of JA responses and a central node integrating multiple hormonal pathways (Song et al., 2022). The observed upregulation of *MYC2* in response to SL deficiency (Figure 5) is a phenomenon already reported in other species including tomato (Xu et al., 2019). Interactions between SL and JA pathways have also been documented (Visentin et al., 2016; Korek and Marzec, 2023; Han et al., 2025). Future studies addressing the contribution of JA and other hormonal signals to the transcriptional activation of MYC2 in SL-deficient backgrounds could help clarify the mechanistic basis of this regulatory connection.

The upregulation of *GXMs* provides a plausible explanation for the hypermethylation of xylan in the *tbl29* background (Grantham et al., 2017; Figure 2b). Elevated MYC2 transcript abundance in the *max3* background suppresses the expression of these *GXM* genes (Figure 3), thereby reducing GlcA methylation. The loss of suppression in the *tbl29 max3 myc2* triple mutant (Figure 4) confirms that MYC2 activity plays an important part in this compensatory mechanism. The presence of canonical MYC2-binding motifs in the promoter regions of all three *GXM* genes raises the possibility that MYC2 directly represses their expression (Table S1). However, indirect regulation through additional transcriptional factors remains equally plausible. MYC2 has previously been shown to interact with broader transcriptional networks controlling secondary wall biosynthesis, including NAC and MYB regulators, suggesting that *GXM* repression could involve multiple layers of transcriptional control (Ramirez et al., 2011; Ramírez et al., 2011; Ko et al., 2012; Didi et al., 2015; Taylor-Teeples et al., 2015; Im and Son, 2025).

Despite these open questions, the unveiled MYC2-dependent regulatory framework provides a molecular link between SL hormonal signalling and the maintenance of wall structural homeostasis. Our study provides the first evidence that strigolactone signalling can influence the methylation pattern of glucuronoxylan and thereby modulate secondary wall composition in response to defects in xylan *O*-acetylation. Beyond its established roles in developmental processes, our findings build upon previous reports pointing to a role of SL in regulating SCW formation (Agusti et al., 2011; Agusti, 2017). The interplay between SL and JA signalling via MYC2 may provide a mechanism by which plants adjust xylan MeGlcA levels in response to perturbations in *O*-acetylation. More broadly, our results suggest that compensatory pathways do not necessarily restore the original wall architecture when cell wall biosynthesis is perturbed, but can instead achieve a functionally equivalent state through remodelling of polysaccharide substitution patterns controlled by hormonal signals. Such plasticity would contribute to developmental robustness and stress resilience by maintaining secondary wall functionality under genetic or environmental challenges.

## EXPERIMENTAL PROCEDURES

### Plant materials and growth conditions

All plants were *Arabidopsis thaliana* Columbia-0 (Col-0) ecotype. The mutant lines *tbl29* (SAIL_856_G11), *max3-9* (N9567), *gux1* (SALK_063763), and *jin1-9*/*myc2* (SALK_017005c) were obtained from the NASC (http://arabidopsis.info) (Scholl et al., 2000) and ABRC (https://abrc.osu.edu) stock centers. Homozygous lines were identified by PCR genotyping using specific primer combinations.

The double mutant *tbl29max3-9* (Ramirez and Pauly, 2019) was crossed with *gux1*, and the F_2_ progeny were screened to obtain the *tbl29max3-9gux1* triple mutant. Similarly, *tbl29max3-9* was crossed with *myc2*, and the resulting F_2_ generation was screened to identify the *tbl29max3-9myc2* triple mutant. Primer sequences used for genotyping all mutants are listed in Table S1.

Seeds were stratified in 0.2% agarose for 4 days at 4 °C in darkness, then sown on soil and grown in a controlled phytotron (23 °C/21 °C, 16 h light/8 h dark, 8,000 lux).

### Cell wall composition analysis

Six-week-old mature primary stems were collected and freeze-dried at –100 °C under vacuum for 2 days, then ground into a fine powder at 30 Hz for 2 min using an MM400 mixer mill (Retsch). Preparation of de-starched alcohol-insoluble residue (dAIR) was performed as described by (Foster et al., 2010).

For the uronic acids composition analysis, 1 mg of dAIR was hydrolyzed in 2 M trifluoroacetic acid (TFA) at 121 °C for 90 min. The hydrolysates were washed twice with 2-propanol, resuspended in deionized water, and determined by high-performance anion-exchange chromatography with pulsed amperometric detection (HPAEC-PAD) (Knauer). The MeGlcA and GlcA were separated using a CarboPac PA200 analytical column (3 x 250 mm) with a CarboPac PA200 guard column (3 x 50 mm) (Thermo Fisher Scientific) at a flow rate of 0.5 mL/min and 40 °C. The eluents were: A: double-distilled H_2_O, B: 200 mM NaOH containing 400 mM NaOAc (Sigma-Aldrich), C: 200 mM NaOH. Separation followed the elongated gradient method described by (Leontakianakou et al., 2024).

### MALDI–TOF MS analysis

For the determination of the [Me]GlcA substituent pattern in different genotypes, 1 mg of dAIR from each sample was deacetylated by incubation with 100 µL of 0.5 M NaOH at 25 °C for 1 h under constant shaking. The reaction was neutralized by adding an equal volume of 1 M HCl, followed by centrifugation to remove the supernatant. The pellet was washed once by resuspension in 1 mL of water and centrifuged again. The resulting deacetylated pellet was resuspended in 99 µL of ddH_2_O, then 1 µL of CjXyn10A (NZYtech) was added. The mixture was incubated at 37 °C for 20 h with constant shaking to perform xylanase digestion. Xylooligosaccharide-containing supernatants were collected by centrifugation, and 1 µL of each sample was spotted onto a MALDI target plate together with 1 µL of matrix solution (2,5-dihydroxybenzoic acid, 10 mg mL^−1^ in 10 mM NaCl). Positive-ion MALDI–TOF mass spectra were acquired using a RapifleX MALDI–TOF (Bruker) instrument operating in reflectron mode with an acceleration voltage of 20 kV. The relative abundance of xylan oligosaccharides with different [Me]GlcA substituents was calculated based on the peak area of the corresponding ions in the MALDI–TOF spectra.

### Gene expression analysis

Primary stems from 6-week-old plants were harvested and immediately ground into fine powder in liquid nitrogen. Total RNA was extracted using the RNeasy Plant Mini Kit (Qiagen) following the manufacturer’s instructions. One microgram of RNA was treated with TURBO DNase I (Thermo Fisher Scientific) to remove genomic DNA contamination, and first-strand cDNA was synthesized using the iScript Reverse Transcription Supermix (Bio-Rad). Quantitative real-time PCR (qRT-PCR) was performed with the resulting cDNA as template using the SsoAdvanced Universal SYBR Green Supermix (Bio-Rad) on a CFX96 Real-Time PCR Detection System (Bio-Rad). Relative gene expression levels were normalized against ACTIN2 (At3g18780) as the reference gene. Primer sequences used for qRT-PCR are provided in Table S2.

### Statistical analysis

All experiments were independently performed at least three biological replicates. Statistical significance among multiple groups was determined by one-way ANOVA followed by Tukey’s multiple comparison test (P < 0.05). Different letters indicate significant differences among groups.

## AUTHOR CONTRIBUTIONS

VR and MP conceived and designed the project. SW performed experiments and analyzed data. SW, MP and VR wrote the manuscript. All authors contributed to the article and approved the submitted version.

## ACKNOWLEDGMENTS

The authors would like to thank Katharina Grosche and Felix Roth for excellent technical support in cell wall analyses and plant molecular biology experiments, respectively. This work was funded by the Cluster of Excellence on Plant Sciences (CEPLAS) by the Deutsche Forschungsgemeinschaft (DFG, German Research Foundation) under Germany’s Excellence Strategy-EXC 2048/1-Project ID: 390686111.

## DATA AVAILABILITY STATEMENT

All data supporting the findings of this study are available within the paper and its Supporting Information.

## SUPPORTING INFORMATION

**Figure S1**. *MALDI-ToF-MS* spectra of deacetylated xylan hydrolysed with GH10 xylanase.

**Figure S2**. *GUX1* shows comparable transcription levels in the *tbl29* and *tbl29* max3 mutants.

**Figure S3**. Morphological phenotypes of *gux1, max3 gux1 and tbl29:GUX1-OE* plants

**Table S1**. Cis-acting regulatory elements identified within the 2-kb upstream regions of *GXM* genes.

**Table S2**. Primers used in this study.

## REFERENCES

Agusti, J. (2017) Strigolactone-mediated Stimulation of Secondary Xylem Proliferation in Stems. Methods in Molecular Biology, 1544, 21–26.

Agusti, J., Herold, S., Schwarz, M., Sanchez, P., Ljung, K., Dun, E.A. et al. (2011) Strigolactone signaling is required for auxin-dependent stimulation of secondary growth in plants. Proceedings of the National Academy of Sciences of the United States of America, 108, 20242–20247.

Anderson, J.P., Badruzsaufari, E., Schenk, P.M., Manners, J.M., Desmond, O.J., Ehlert, C. et al. (2004) Antagonistic interaction between abscisic acid and jasmonate-ethylene signaling pathways modulates defense gene expression and disease resistance in Arabidopsis. Plant Cell, 16, 3460–3479.

Bensussan, M., Lefebvre, V., Ducamp, A., Trouverie, J., Gineau, E., Fortabat, M.N. et al. (2015) Suppression of Dwarf and irregular xylem Phenotypes Generates Low-Acetylated Biomass Lines in Arabidopsis. Plant Physiology, 168, 452–463.

Booker, J., Auldridge, M., Wills, S., McCarty, D., Klee, H. and Leyser, O. (2004) MAX3/CCD7 is a carotenoid cleavage dioxygenase required for the synthesis of a novel plant signaling molecule. Current Biology, 14, 1232–1238.

Bouchabke-Coussa, O., Quashie, M.L., Seoane-Redondo, J., Fortabat, M.N., Gery, C., Yu, A. et al. (2008) ESKIMO1 is a key gene involved in water economy as well as cold acclimation and salt tolerance. BMC Plant Biology, 8, 125.

Bromley, J.R., Busse-Wicher, M., Tryfona, T., Mortimer, J.C., Zhang, Z., Brown, D.M. et al. (2013) GUX1 and GUX2 glucuronyltransferases decorate distinct domains of glucuronoxylan with different substitution patterns. The Plant Journal, 74, 423–434.

Delmer, D., Dixon, R.A., Keegstra, K. and Mohnen, D. (2024) The plant cell wall-dynamic, strong, and adaptable-is a natural shapeshifter. Plant Cell, 36, 1257–1311.

Didi, V., Jackson, P. and Hejátko, J. (2015) Hormonal regulation of secondary cell wall formation. Journal of Experimental Botany, 66, 5015–5027.

Escudero, V., Jorda, L., Sopena-Torres, S., Melida, H., Miedes, E., Munoz-Barrios, A. et al. (2017) Alteration of cell wall xylan acetylation triggers defense responses that counterbalance the immune deficiencies of plants impaired in the beta-subunit of the heterotrimeric G-protein. The Plant Journal, 92, 386–399.

Foster, C.E., Martin, T.M. and Pauly, M. (2010) Comprehensive compositional analysis of plant cell walls (lignocellulosic biomass) part II: carbohydrates. Journal of visualized experiments: JoVE, 1837.

Gao, Y., He, C., Zhang, D., Liu, X., Xu, Z., Tian, Y. et al. (2017) Two Trichome Birefringence-Like Proteins Mediate Xylan Acetylation, Which Is Essential for Leaf Blight Resistance in Rice. Plant Physiology, 173, 470–481.

Gille, S. and Pauly, M. (2012) O-acetylation of plant cell wall polysaccharides. Frontiers in Plant Science, 3, 12.

Grantham, N.J., Wurman-Rodrich, J., Terrett, O.M., Lyczakowski, J.J., Stott, K., Iuga, D. et al. (2017) An even pattern of xylan substitution is critical for interaction with cellulose in plant cell walls. Nature Plants, 3, 859–865.

Han, Y., Sun, Y., Wang, H., Li, H., Jiang, M., Liu, X. et al. (2025) Biosynthesis and Signaling of Strigolactones Act Synergistically With That of ABA and JA to Enhance Verticillium dahliae Resistance in Cotton (Gossypium hirsutum L.). Plant, Cell & Environment, 48, 571–586.

Im, J.H. and Son, S. (2025) MYC2 signaling in secondary cell wall modulation. Frontiers in Plant Science, 16, 1558922.

Im, J.H., Son, S., Kim, W.C., Kim, K., Mitsuda, N., Ko, J.H. et al. (2024) Jasmonate activates secondary cell wall biosynthesis through MYC2-MYB46 module. The Plant Journal, 117, 1099–1114.

Joshi, A. and Gupta, M. (2026) Regulatory role of O3 acetylation as a molecular switch in xylan-cellulose interactions. Cellulose, 33, 1965–1983.

Jung, C., Zhao, P., Seo, J.S., Mitsuda, N., Deng, S. and Chua, N.H. (2015) PLANT U-BOX PROTEIN10 Regulates MYC2 Stability in Arabidopsis. Plant Cell, 27, 2016–2031.

Kang, X., Kirui, A., Dickwella Widanage, M.C., Mentink-Vigier, F., Cosgrove, D.J. and Wang, T. (2019) Lignin-polysaccharide interactions in plant secondary cell walls revealed by solid-state NMR. Nature Communications, 10, 347.

Kirui, A., Zhao, W., Deligey, F., Yang, H., Kang, X., Mentink-Vigier, F. et al. (2022) Carbohydrate-aromatic interface and molecular architecture of lignocellulose. Nature Communications, 13, 538.

Ko, J.-H., Kim, W.-C., Kim, J.-Y., Ahn, S.-J. and Han, K.-H. (2012) MYB46-Mediated Transcriptional Regulation of Secondary Wall Biosynthesis. Molecular Plant, 5, 961–963.

Korek, M. and Marzec, M. (2023) Strigolactones and abscisic acid interactions affect plant development and response to abiotic stresses. BMC Plant Biology, 23, 314.

Kundu, T., Smith, J.C. and Gupta, M. (2025) Effect of Acetylation Patterns of Xylan on Interactions with Cellulose. Biomacromolecules, 26, 1659–1671.

Lee, C., Teng, Q., Zhong, R. and Ye, Z.H. (2011) The four Arabidopsis reduced wall acetylation genes are expressed in secondary wall-containing cells and required for the acetylation of xylan. Plant Cell Physiology, 52, 1289–1301.

Lee, C., Teng, Q., Zhong, R. and Ye, Z.H. (2012a) Arabidopsis GUX proteins are glucuronyltransferases responsible for the addition of glucuronic acid side chains onto xylan. Plant Cell Physiology, 53, 1204–1216.

Lee, C., Teng, Q., Zhong, R., Yuan, Y., Haghighat, M. and Ye, Z.H. (2012b) Three Arabidopsis DUF579 domain-containing GXM proteins are methyltransferases catalyzing 4-o-methylation of glucuronic acid on xylan. Plant Cell Physiology, 53, 1934–1949.

Lefebvre, V., Fortabat, M.N., Ducamp, A., North, H.M., Maia-Grondard, A., Trouverie, J. et al. (2011) ESKIMO1 disruption in Arabidopsis alters vascular tissue and impairs water transport. PLoS One, 6, e16645.

Leontakianakou, S., Grey, C., Karlsson, E.N. and Sardari, R.R. (2024) An improved HPAEC-PAD method for the determination of D-glucuronic acid and 4-O-methyl-D-glucuronic acid from polymeric and oligomeric xylan. BMC Biotechnology, 24, 100.

Lescot, M., Déhais, P., Thijs, G., Marchal, K., Moreau, Y., Van de Peer, Y. et al. (2002) PlantCARE, a database of plant cis-acting regulatory elements and a portal to tools for in silico analysis of promoter sequences. Nucleic Acids Research, 30, 325–327.

Li, R., Weldegergis, B.T., Li, J., Jung, C., Qu, J., Sun, Y. et al. (2014) Virulence Factors of Geminivirus Interact with MYC2 to Subvert Plant Resistance and Promote Vector Performance Plant Cell, 26, 4991–5008.

Lopez-Vidriero, I., Godoy, M., Grau, J., Penuelas, M., Solano, R. and Franco-Zorrilla, J.M. (2021) DNA features beyond the transcription factor binding site specify target recognition by plant MYC2-related bHLH proteins. Plant Communications, 2, 100232.

Luo, F., Zhang, Q., Xin, H., Liu, H., Yang, H., Doblin, M.S. et al. (2022) A Phytochrome B-PIF4-MYC2/MYC4 module inhibits secondary cell wall thickening in response to shaded light. Plant Communications, 3, 100416.

Manabe, Y., Verhertbruggen, Y., Gille, S., Harholt, J., Chong, S.L., Pawar, P.M. et al. (2013) Reduced Wall Acetylation proteins play vital and distinct roles in cell wall O-acetylation in Arabidopsis. Plant Physiology, 163, 1107–1117.

Martinez-Abad, A., Berglund, J., Toriz, G., Gatenholm, P., Henriksson, G., Lindstrom, M. et al. (2017) Regular Motifs in Xylan Modulate Molecular Flexibility and Interactions with Cellulose Surfaces. Plant Physiology, 175, 1579–1592.

Mortimer, J.C., Miles, G.P., Brown, D.M., Zhang, Z., Segura, M.P., Weimar, T. et al. (2010) Absence of branches from xylan in Arabidopsis gux mutants reveals potential for simplification of lignocellulosic biomass. Proceedings of the National Academy of Sciences of the United States of America, 107, 17409–17414.

Nakano, Y., Yamaguchi, M., Endo, H., Rejab, N.A. and Ohtani, M. (2015) NAC-MYB-based transcriptional regulation of secondary cell wall biosynthesis in land plants. Frontiers in Plant Science, 6, 288.

Pauly, M., Gille, S., Liu, L., Mansoori, N., de Souza, A., Schultink, A. et al. (2013) Hemicellulose biosynthesis. Planta, 238, 627–642.

Pauly, M. and Ramirez, V. (2018) New Insights Into Wall Polysaccharide O-Acetylation. Frontiers in Plant Science, 9, 1210.

Pereira, C.S., Silveira, R.L., Dupree, P. and Skaf, M.S. (2017) Effects of Xylan Side-Chain Substitutions on Xylan-Cellulose Interactions and Implications for Thermal Pretreatment of Cellulosic Biomass. Biomacromolecules, 18, 1311–1321.

Pfaff, S.A., Wagner, E.R. and Cosgrove, D.J. (2024) The structure and interaction of polymers affects secondary cell wall banding patterns in Arabidopsis. Plant Cell, 36, 4309–4322.

Qaseem, M.F., Zhang, W., Dupree, P. and Wu, A.M. (2025) Xylan structural diversity, biosynthesis, and functional regulation in plants. The International Journal of Biological Macromolecules, 291, 138866.

Ramirez, V., Agorio, A., Coego, A., Garcia-Andrade, J., Hernandez, M.J., Balaguer, B. et al. (2011) MYB46 modulates disease susceptibility to Botrytis cinerea in Arabidopsis. Plant Physiology, 155, 1920–1935.

Ramírez, V., García-Andrade, J. and Vera, P. (2011) Enhanced disease resistance to Botrytis cinerea in myb46 Arabidopsis plants is associated to an early down-regulation of CesA genes. Plant Signaling & Behavior, 6, 911–913.

Ramirez, V. and Pauly, M. (2019) Genetic dissection of cell wall defects and the strigolactone pathway in Arabidopsis. Plant Direct, 3, e00149.

Ramirez, V., Xiong, G., Mashiguchi, K., Yamaguchi, S. and Pauly, M. (2018) Growth- and stress-related defects associated with wall hypoacetylation are strigolactone-dependent. Plant Direct, 2, e00062.

Scheller, H.V. and Ulvskov, P. (2010) Hemicelluloses. Annual Review of Plant Biology, 61, 263–289.

Scholl, R.L., May, S.T. and Ware, D.H. (2000) Seed and molecular resources for Arabidopsis. Plant Physiology, 124, 1477–1480.

Schultink, A., Naylor, D., Dama, M. and Pauly, M. (2015) The role of the plant-specific ALTERED XYLOGLUCAN9 protein in Arabidopsis cell wall polysaccharide O-acetylation. Plant Physiology, 167, 1271–1283.

Simmons, T.J., Mortimer, J.C., Bernardinelli, O.D., Poppler, A.C., Brown, S.P., deAzevedo, E.R. et al. (2016) Folding of xylan onto cellulose fibrils in plant cell walls revealed by solid-state NMR. Nature Communications, 7, 13902.

Song, C., Cao, Y., Dai, J., Li, G., Manzoor, M.A., Chen, C. et al. (2022) The Multifaceted Roles of MYC2 in Plants: Toward Transcriptional Reprogramming and Stress Tolerance by Jasmonate Signaling. Frontiers in Plant Science, 13, 868874.

Sorefan, K., Booker, J., Haurogne, K., Goussot, M., Bainbridge, K., Foo, E. et al. (2003) MAX4 and RMS1 are orthologous dioxygenase-like genes that regulate shoot branching in Arabidopsis and pea. Genes & Development 17, 1469–1474.

Sun, A., Yu, B., Zhang, Q., Peng, Y., Yang, J., Sun, Y. et al. (2020) MYC2-Activated TRICHOME BIREFRINGENCE-LIKE37 Acetylates Cell Walls and Enhances Herbivore Resistance. Plant Physiology, 184, 1083–1096.

Taylor-Teeples, M., Lin, L., de Lucas, M., Turco, G., Toal, T.W., Gaudinier, A. et al. (2015) An Arabidopsis gene regulatory network for secondary cell wall synthesis. Nature, 517, 571–575.

Tryfona, T., Bourdon, M., Delgado Marques, R., Busse-Wicher, M., Vilaplana, F., Stott, K. et al. (2023) Grass xylan structural variation suggests functional specialization and distinctive interaction with cellulose and lignin. The Plant Journal, 113, 1004–1020.

Urbanowicz, B.R., Pena, M.J., Ratnaparkhe, S., Avci, U., Backe, J., Steet, H.F. et al. (2012) 4-O-methylation of glucuronic acid in Arabidopsis glucuronoxylan is catalyzed by a domain of unknown function family 579 protein. Proceedings of the National Academy of Sciences of the United States of America, 109, 14253–14258.

Visentin, I., Vitali, M., Ferrero, M., Zhang, Y., Ruyter-Spira, C., Novák, O. et al. (2016) Low levels of strigolactones in roots as a component of the systemic signal of drought stress in tomato. New Phytologist, 212, 954–963.

Xin, Z. and Browse, J. (1998) eskimo1 mutants of Arabidopsis are constitutively freezing-tolerant. Proceedings of the National Academy of Sciences of the United States of America, 95, 7799–7804.

Xiong, G., Cheng, K. and Pauly, M. (2013) Xylan O-acetylation impacts xylem development and enzymatic recalcitrance as indicated by the Arabidopsis mutant tbl29. Molecular Plant, 6, 1373–1375.

Xiong, G., Dama, M. and Pauly, M. (2015) Glucuronic Acid Moieties on Xylan Are Functionally Equivalent to O-Acetyl-Substituents. Molecular Plant, 8, 1119–1121.

Xu, X., Fang, P., Zhang, H., Chi, C., Song, L., Xia, X. et al. (2019) Strigolactones positively regulate defense against root-knot nematodes in tomato. Journal of Experimental Botany, 70, 1325–1337.

Ye, Z.H. and Zhong, R. (2022) Outstanding questions on xylan biosynthesis. Plant Science, 325, 111476.

Yuan, Y., Teng, Q., Lee, C., Zhong, R. and Ye, Z.H. (2014) Modification of the degree of 4-O-methylation of secondary wall glucuronoxylan. Plant Science, 219-220, 42–50.

Yuan, Y., Teng, Q., Zhong, R. and Ye, Z.H. (2013) The Arabidopsis DUF231 domain-containing protein ESK1 mediates 2-O- and 3-O-acetylation of xylosyl residues in xylan. Plant Cell Physiology, 54, 1186–1199.

Yuan, Y., Teng, Q., Zhong, R. and Ye, Z.H. (2016a) Roles of Arabidopsis TBL34 and TBL35 in xylan acetylation and plant growth. Plant Science, 243, 120–130.

Yuan, Y., Teng, Q., Zhong, R. and Ye, Z.H. (2016b) TBL3 and TBL31, Two Arabidopsis DUF231 Domain Proteins, are Required for 3-O-Monoacetylation of Xylan. Plant Cell Physiology, 57, 35–45.

Zhang, B., Gao, Y., Zhang, L. and Zhou, Y. (2021) The plant cell wall: Biosynthesis, construction, and functions. Journal of Integrative Plant Biology, 63, 251–272.

Zhang, J., Chen, W., Li, X., Shi, H., Lv, M., He, L. et al. (2023) Jasmonates regulate apical hook development by repressing brassinosteroid biosynthesis and signaling. Plant Physiology, 193, 1561–1579.

Zhong, R., Cui, D. and Ye, Z.H. (2019) Secondary cell wall biosynthesis. New Phytologist, 221, 1703–1723.

Zhong, R. and Ye, Z.-H. (2015) Secondary Cell Walls: Biosynthesis, Patterned Deposition and Transcriptional Regulation. Plant and Cell Physiology, 56, 195–214.

