## Supplemental Figures 1-3 for "MYC2 mediated regulation of xylan substitution patterns"

### Slide 1
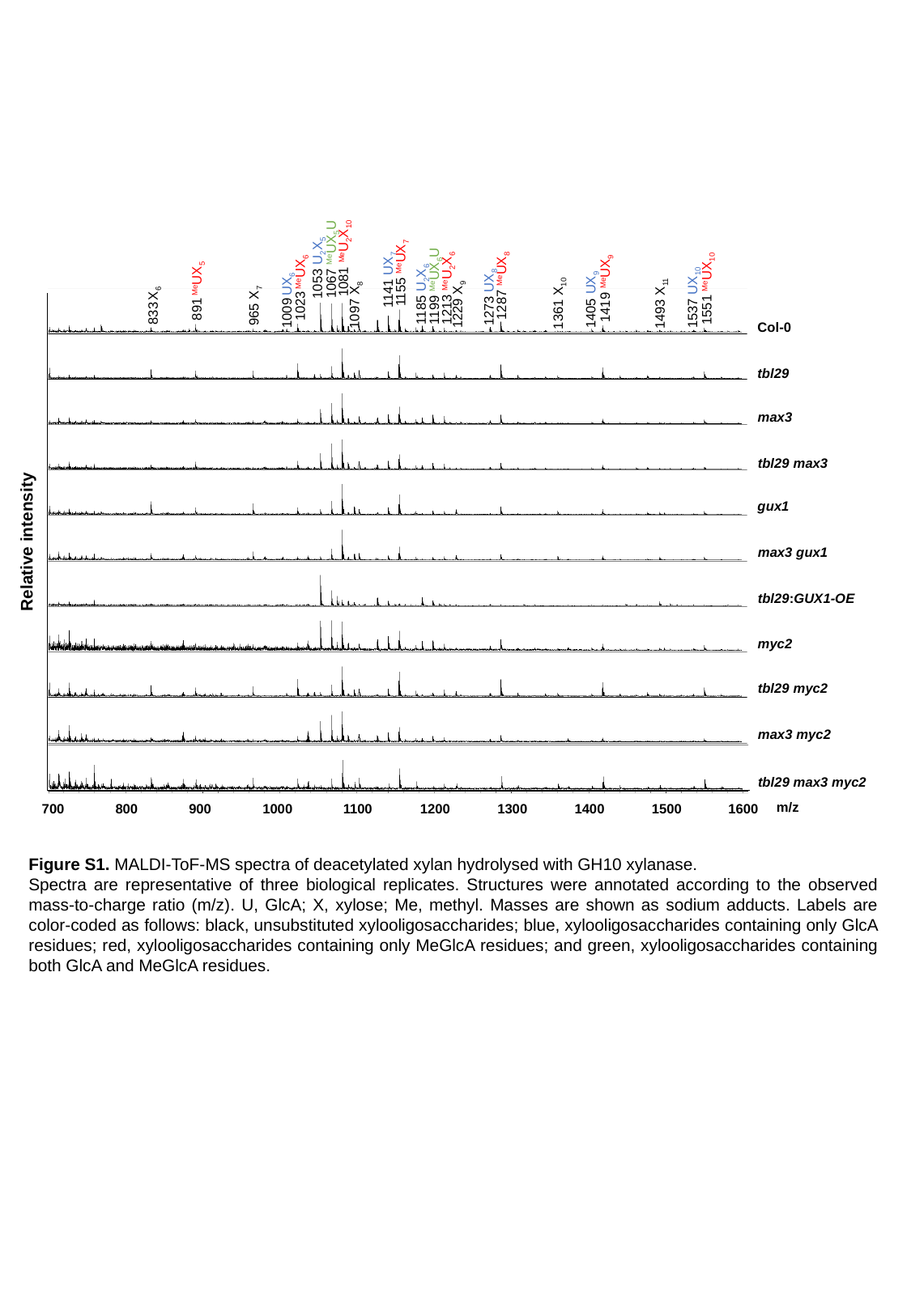

1081 ^Me^U_2_X_10_
1067 ^Me^UX_5_U
1053 U_2_X_5_
1155 ^Me^UX_7_
1141 UX_7_
1287 ^Me^UX_8_
1273 UX_8_
1419 ^Me^UX_9_
1405 UX_9_
1199 ^Me^UX_6_U
1023 ^Me^UX_6_
1213 ^Me^U_2_X_6_
1551 ^Me^UX_10_
891 ^Me^UX_5_
1185 U_2_X_6_
1537 UX_10_
1009 UX_6_
Col-0
tbl29
max3
tbl29 max3
gux1
Relative intensity
max3 gux1
tbl29:GUX1-OE
myc2
tbl29 myc2
max3 myc2
tbl29 max3 myc2
m/z
700
800
900
1000
1100
1200
1300
1400
1500
1600
1361 X_10_
1493 X_11_
1229 X_9_
1097 X_8_
833 X_6_
965 X_7_
Figure S1. MALDI-ToF-MS spectra of deacetylated xylan hydrolysed with GH10 xylanase.
Spectra are representative of three biological replicates. Structures were annotated according to the observed mass-to-charge ratio (m/z). U, GlcA; X, xylose; Me, methyl. Masses are shown as sodium adducts. Labels are color-coded as follows: black, unsubstituted xylooligosaccharides; blue, xylooligosaccharides containing only GlcA residues; red, xylooligosaccharides containing only MeGlcA residues; and green, xylooligosaccharides containing both GlcA and MeGlcA residues.

### Slide 2
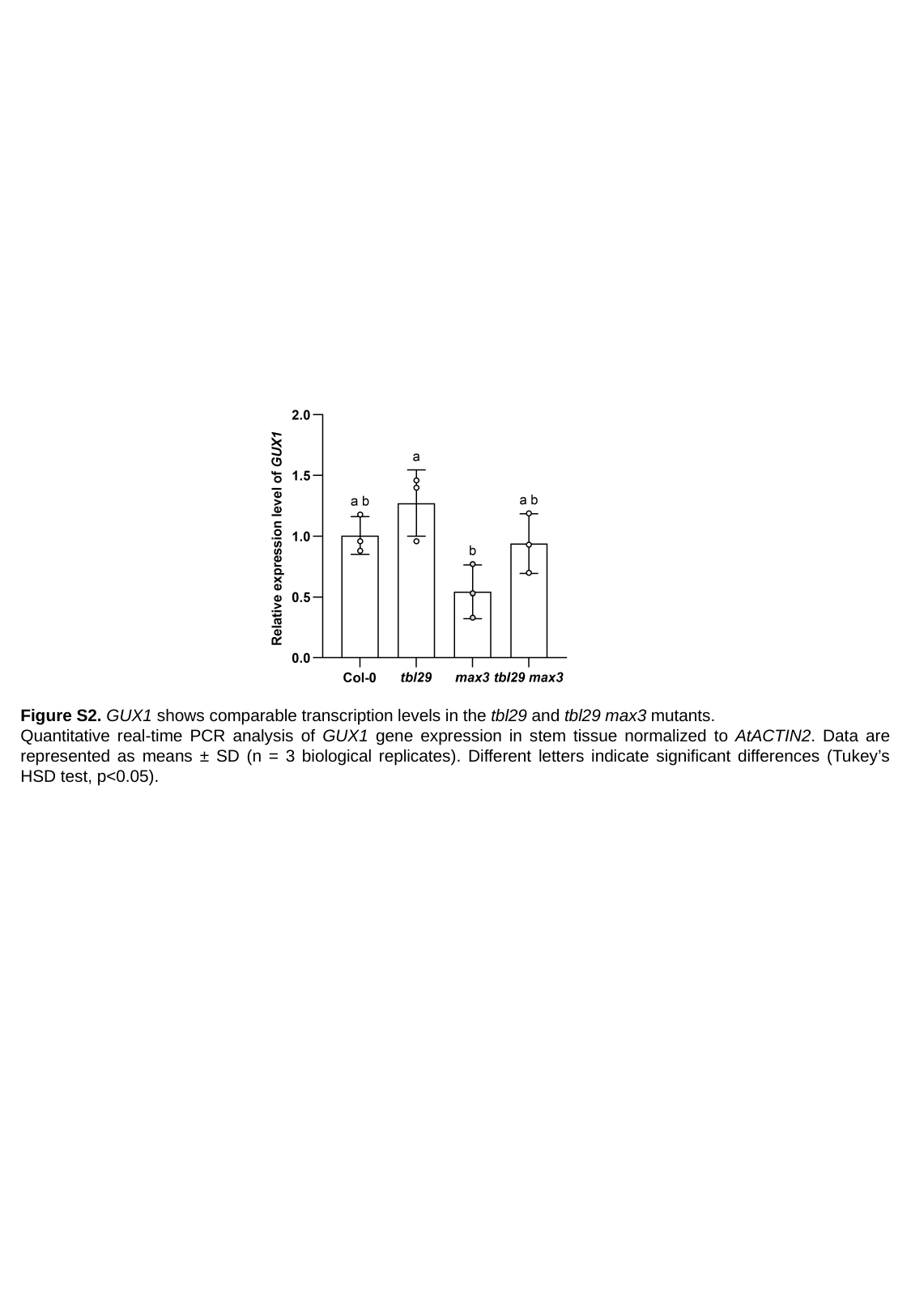

Figure S2. GUX1 shows comparable transcription levels in the tbl29 and tbl29 max3 mutants.
Quantitative real-time PCR analysis of GUX1 gene expression in stem tissue normalized to AtACTIN2. Data are represented as means ± SD (n = 3 biological replicates). Different letters indicate significant differences (Tukey’s HSD test, p<0.05).

### Slide 3
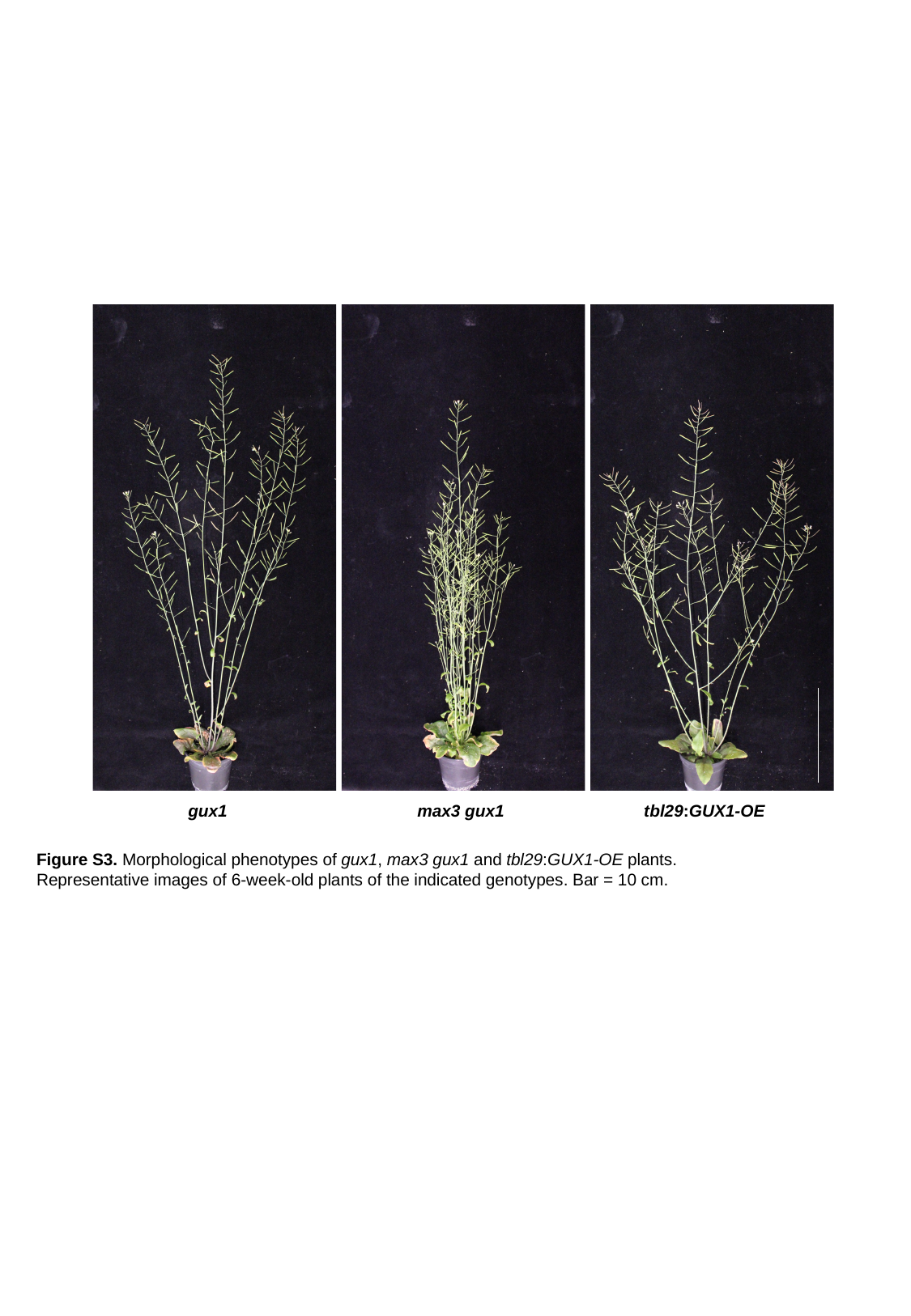

gux1
max3 gux1
tbl29:GUX1-OE
Figure S3. Morphological phenotypes of gux1, max3 gux1 and tbl29:GUX1-OE plants.
Representative images of 6-week-old plants of the indicated genotypes. Bar = 10 cm.
