## Supplemental Table 2 for "MYC2 mediated regulation of xylan substitution patterns"

**Table S2.** List of primers used in this study.

| Primer name | Sequence (5’-3’) |
| --- | --- |
| tbl29-F | AATTTGCAAGCAAAGCATCAC |
| tbl29-R | TGGGTTTTTGATAACGAGACG |
| LB1 | GCCTTTTCAGAAATGGATAAATAGCCTTGCTTCC |
| max3-9-F | AGGTGTATTTAAGATGCCA |
| max3-9-R | CACAAAATGTGAAGTTGCTT |
| myc2-F | GGCGGGATTTAATCAAGAGAC |
| myc2-R | TTTGGTACAACCGCTCGTAAC |
| gux1-F | ATCTCCGGTCTACGGTAATGG |
| gux1-R | CTACAGCAAGTTCCGGCTATG |
| LBb1.3 | ATTTTGCCGATTTCGGAAC |
| AtACTIN2-qRT-F | TCTTCCGCTCTTTCTTTCCAAGC |
| AtACTIN2-qRT-R | ACCATTGTCACACACGATTGGTTG |
| MYC2-qRT-F | TATGTCGGTGGTTAACGATTTG |
| MYC2-qRT-R | TCCCAAAACACACCCTTTTAAC |
| GXM1-qRT-F | ACAGGAAACAGGGAGGAGAGACTG |
| GXM1-qRT-R | CTATGGACAAAAGGGTCTATTTGATTC |
| GXM2-qRT-F | TTGGCTCGTAACCGTTATGACGGT |
| GXM2-qRT-R | TCAAAAGCGGCGACTGATATCCGC |
| GXM3-qRT-F | TGGCTCGTAACCGTGAAGATGGTG |
| GXM3-qRT-R | TTAACGGCGGCGATCAACTTCCAC |
